# Allosteric site variants affect GTP hydrolysis on RAS

**DOI:** 10.1101/2023.05.29.542740

**Authors:** Christian W. Johnson, Susan K. Fetics, Kathleen P. Davis, Jose A. Rodrigues, Carla Mattos

**Author notes:** Please address all correspondence to Dr. Carla Mattos at. Description of supplementary material: 2 tables describing crystal statistics, 1 table describing crystallization conditions, and 2 tables describing rates and rate constants for GTP hydrolysis reactions by H-Ras R97 mutants. **Impact statement**: Small GTPases regulate cell signaling by switching between GTP– and GDP-bound states. Transition to the GDP-bound state depends on GTP hydrolysis. Here, we show that GTP hydrolysis in HRAS is altered by mutations outside its active site, providing a rare demonstration of allostery in a small GTPase. This allosteric function may relate to RAS interaction with RAF kinases and reflects an underappreciated oncogenic mechanism of HRAS, NRAS, and KRAS.

## Abstract

RAS GTPases are proto-oncoproteins that regulate cell growth, proliferation, and differentiation in response to extracellular signals. The signaling functions of RAS, and other small GTPases, are dependent on their ability to cycle between GDP-bound and GTP-bound states. Structural analyses suggest that GTP hydrolysis catalyzed by HRAS can be regulated by an allosteric site located between helices 3, 4 and loop 7. Here we explore the relationship between intrinsic GTP hydrolysis on HRAS and the position of helix 3 and loop 7 through manipulation of the allosteric site, showing that the two sites are functionally connected. We generated several hydrophobic mutations in the allosteric site of HRAS to promote shifts in helix 3 relative to helix 4. By combining crystallography and enzymology to study these mutants, we show that closure of the allosteric site correlates with increased hydrolysis of GTP on HRAS in solution. Interestingly, binding to the RAS binding domain of RAF kinase (RAF-RBD) inhibits GTP hydrolysis in the mutants. This behavior may be representative of a cluster of poorly understood mutations that occur in human tumors, which potentially cooperate with RAF complex formation to stabilize the GTP-bound state of RAS.

## Introduction

Understanding how proto-oncogenes regulate cell signaling is critical to understanding their role in both homeostasis and disease. For instance, KRAS, NRAS, and HRAS (collectively RAS) are peripheral membrane hub proteins that activate many different signaling pathways, including the RAF/MEK/ERK signaling pathway, which is essential for cell differentiation, proliferation and survival ^1^. RAS proteins regulate cell signaling through the controlled cycling of GTP– and GDP-bound states. When bound to GTP, RAS is in a signaling active state that interacts with downstream effectors, such as RAF kinase ^2;3^. When Ras is bound to GDP, it is in its inactive state that cannot bind effectors. Point-mutations in KRAS, NRAS and HRAS act to stabilize the GTP bound state and are found in 16%, 2.6% and 0.9% of all cancers, respectively ^4^. Unfortunately, despite the recent approval of KRAS-specific inhibitors Sotorasib and Adagrasib which target G12C mutations, the vast majority of KRAS, NRAS, and HRAS mutants remain elusive to therapeutic intervention ^5^. Therefore, more work is needed to understand the molecular mechanisms through which RAS and its effectors work.

To maintain homeostasis, inactivation of RAS is driven by hydrolysis of GTP to GDP. This can occur through GTPase-activating proteins (GAPs) which bind to the active site of RAS ^6^. This mechanism is widely viewed as critical to maintaining the homeostatic functions of RAS, because many GAPs (e.g. NF1, p120GAP) are found inactivated in tumors ^4^. However, regulation of the GTP bound state of RAS by GAPs alone is difficult to reconcile for several reasons. The p120GAP (also called RASA1) protein cannot compete with Ras effectors, including RAF, under physiological-like conditions ^7^, and it is well documented that RAF blocks GAP activity by binding to RAS ^7-11^. Furthermore, the spectrum of cancers that frequently show NF1 and p120GAP inactivation weakly overlap with those with activating RAS mutations ^4^. Thus, an alternative means to inactivate RAS may rely on intrinsic hydrolysis in the absence of GAP proteins. While this mechanism of hydrolysis occurs slowly under most studied conditions (0.006-0.019 min^-1^) ^12^, we put forth a model whereby intrinsic hydrolysis could be enhanced in a regulated manner from an allosteric site via a network of water-mediated H-bonding interactions to the Ras active site ^13;14^. In this model (**Figure 1A, left**), stabilization of the active site into a closed conformation (e.g. state 2) by RAF kinase, or another effector, increases dependency of the intrinsic hydrolysis reaction on the conformation of switch II (**Figure 1A, right**). Structural analysis suggests that movement of helix 3 toward a pocket on the other side of the protein (**allosteric site,** **Figure 1A**) allows switch II to become ordered in the active site, placing catalytic residues near GTP. We hypothesize that these changes in conformation are responsible for intrinsic GTP hydrolysis.

**Figure 1.**
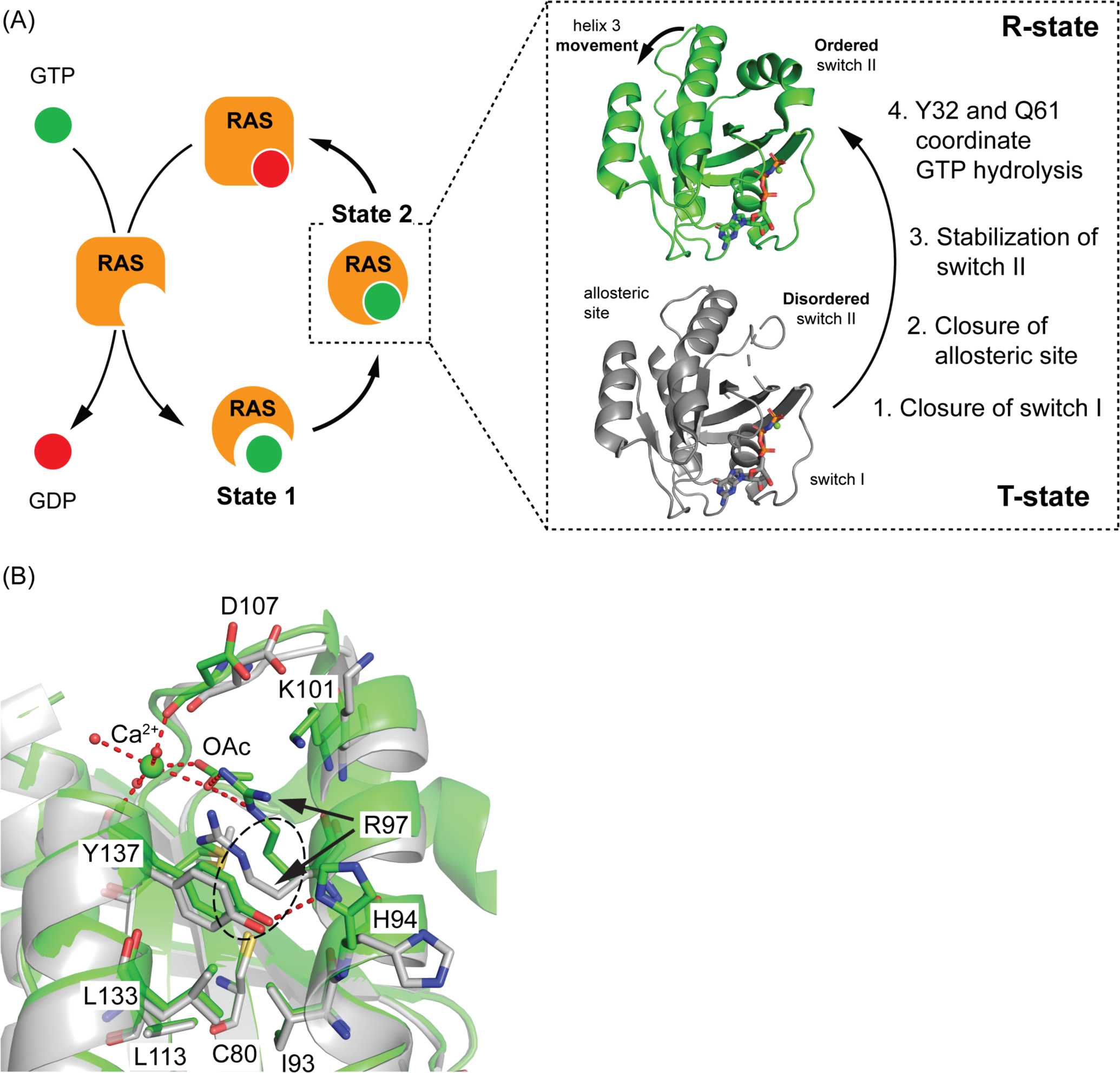
Mutations at R97 induce increased packing in the allosteric site. (A) Model for the regulation of intrinsic hydrolysis by allosteric site transitions. (B) Residue interactions of the allosteric site when HRAS is in the R– and T-states.

The active site of HRAS undergoes several conformational transitions once it becomes bound to GTP ^3;15^. Here, we differentiate the sub-states of state 2 relevant to our model of hydrolysis as the inactive ‘T-state’ (**grey,** **Figure 1A**) and the catalytically active ‘R-state’ (**green,** **Figure 1A**) ^2^. In our previous crystallography experiments, the transition of HRAS from the T-state to the R-state was accomplished by either growing or soaking protein crystals with Ca(OAc)_2_. Calcium and acetate are likely ligand mimetics as they bind weakly to the allosteric site in solution ^16^. In the presence of bound Ca^2+^ and acetate, helix 3 (residues 93-104) and loop 7 (residues 105-108) shift toward helix 4, allowing switch II (residues 60-76) to stabilize the R-state conformation (**green,** **Figure 1B**). This promotes a pre-transition state where Y32 from switch I (residues 28-40) and Q61 from switch II (residues 60-76) interact with a ‘bridging’ water molecule that also forms an H-bond with the γ-phosphate of GTP. This water molecule, which carries a partial positive charge, neutralizes the accumulation of negative charge at the β– and γ-phosphates of GTP during the hydrolysis reaction ^17^. The bridging water molecule is thus positioned to lower the transition state energy associated with GTP hydrolysis similar to the arginine finger introduced by GAPs ^13;18^.

In the present work, we explicitly studied the relationship between allosteric and active sites to understand the mechanism through which the disorder to order transition in switch II completes the active site for GTP hydrolysis. We designed a series of hydrophobic mutants of R97 on HRAS, determined their crystal structures, which revealed their conformation in either R– or T-states, and measured the hydrolysis rate constant for each mutant in the absence and presence of Raf-RBD. In the wild type T-state, R97 is partially buried in a relatively large hydrophobic pocket at the base of the allosteric site (**dotted oval,** **Figure 1B**). In the wild type R-state, R97 transitions out of the hydrophobic pocket and forms a salt-bridge with the acetate molecule, which also interacts with the bound calcium ion. With the help of an H-bond between Y137 (helix 4) and H94 (helix3), the bound calcium and acetate ions promote a shift in helix 3 toward helix 4 (**Figure 1B**). We hypothesized that hydrophobic mutants of R97 would similarly promote the shift of helix 3 toward helix 4, by packing in the hydrophobic pocket, thereby facilitating the R-state transition in a ligand-independent manner.

To test the relationship between the R-state and intrinsic GTP hydrolysis, we crystallized seven R97 mutants in different crystal forms to determine the extent to which each mutation promoted conformations associated with the R-state, and then correlated these changes to measurements of GTP hydrolysis rate constants. By combining our structure and hydrolysis data, we show that closure of the allosteric site (e.g. shift of helix 3 toward helix 4) correlates with an increase in GTP hydrolysis. Unexpectedly, RAF inhibits hydrolysis to varying degrees in a mutation-specific manner. For instance, R97M has a larger hydrolysis rate constant than wild type HRAS, yet unlike the wildtype protein, it shows a ∼50% reduction in activity in the presence of RAF1-RBD. Our data show that some hydrophobic mutations in the allosteric site can enhance the GTP-bound state of RAS when RAF kinases are present, and explain the presence of allosteric site mutations of KRAS found in some human tumors. Furthermore, our work is a proof-of-principle demonstration showing that manipulation of the allosteric site leads to changes in hydrolysis, paving the way for the development of pro–hydrolysis RAS inhibitors.

## Results and Discussion

### R– and T-state classification of crystalized HRAS R97 mutants

HRAS crystals with R32 space group symmetry have switch II and the C-terminal end of helix 3 away from crystal contacts, allowing for transition between R– and T-states in the crystals ^13;19^. Thus, this crystal form is ideal for assessing the effects of the R97 mutants on the switch II/helix 3/allosteric site conformational states (**Supplementary Table 1**). Mutants of HRAS in the R-state showed stabilization of the bridging water molecule in the active site, stabilization of switch II into an ordered helix, and movement of helix 3 toward helix 4 (**Figure 2A**). The nucleophilic water was present in each of our crystal structures (**Figure 2A-B**).

**Figure 2:**
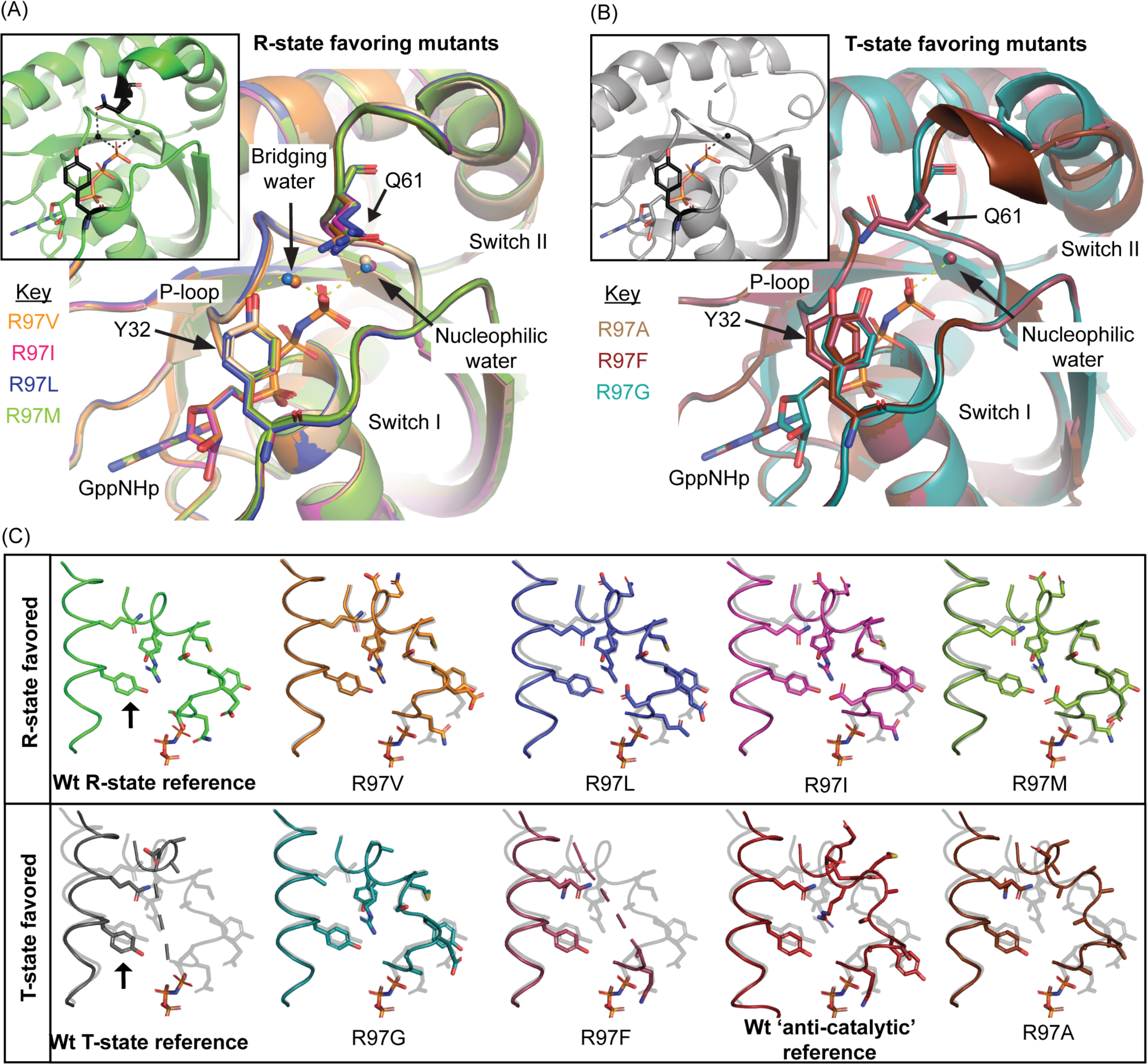
Classification of R32 protein crystals from HRAS R97 mutants. (A) R97V, R97L, R97M, and R97I conserve all features of the R-state except the conformation of Q61. Inset shows wildtype HRAS in the R-state (PDB code 3K8Y ^13^). The catalytic residues Y32 and Q61, and the bridging and nucleophilic waters of wildtype HRAS are shown in black for reference. (B) R97A, R97G, and R97F have active site features similar to the T-state. Inset shows Q61 and nucleophilic water in black for wildtype HRAS in the T-state (PDB code 2RGE^21^). (C) Comparison helix 3 and switch II in each of the R97 mutants. Wildtype reference structures are included in bold for the R-state, T-state and the T-state when HRAS is in the ‘anti-catalytic’ conformation (PDB code 4DLZ ^19^). In light grey is an overlay of the R-state. The black arrow denotes the location of Y96.

The R-state favoring mutants R97V, R97I, R97L, and R97M, stabilized the bridging water molecule in the active site via an H-bond with Y32 (**Figure 2A, yellow dashes**). Likewise, switch II of these mutants was well ordered and formed an H-bond network with helix 3 consistent with the network described for wildtype HRAS in the R-state ^13^, including participation of Y96 which was shifted toward switch II in these mutants (**arrow**, **Figure 2C**). However, despite the overall features of the R-state, none of the R97V, R97I, R97L nor R97M structures captured Q61 in a conformation that allowed H– bonding with the bridging water molecule (**Figure 2A**).

The R97A, R97F, and R97G structures showed active site and switch II features more like the T-state, including absence of the bridging water molecule and a direct interaction between the side chain of Y32 and the γ-phosphate of GTP (**Figure 2B**). Typically switch II in the T-state is disordered (residues 61-69), however it can be stabilized in an ‘anti-catalytic’ conformation that is overall similar to small GTPase RAN in complex with IMPORTIN-β (PDB code 1IBR ^20^), which we previously also observed for the oncogenic mutant HRAS Q61L ^13;21^. While R97A adopts the ‘anti-catalytic’ conformation with helix 3 in the T-state, R97F is in the T-state with an extended disorder of switch II, from residues 61 to 72 (**Figure 2C**). In R97G, electron density supports a switch II helical structure similar to the R-state (**Figure 2C**), although a second conformation of key residues associated with the T-state is also supported by the electron density maps (*not shown*). Consistent with these features, Y96 in R97G is in an R-state position, whereas Y96 in R97A and R97F are in a T-state position (**Figure 2C**).

The overall structural features observed for the allosteric site mutants support our initial hypothesis that mutation of R97 to hydrophobic residues with side chains of moderate size induces a shift of helix 3 to promote the R-state independent of bound Ca^2+^ and acetate. This is consistent with our expectation of a functional connection between the allosteric and active sites of HRAS. As shown in **Supplementary Table 2**, there was no apparent correlation between crystallization condition (i.e. Ca(AOc)_2_, PEG) and whether the crystallized proteins favored the R– or T-states, indicating that the R97 mutation is the main determinant. However, given that only a subset of these mutations induces this shift suggests that both the extent of side chain packing, and the relative position of those amino acid residues in the packing interaction, play a role in the transitions between R– and T-states. This is exemplified by our R97A, R97F and R97G mutants. R97F appears too large to pack in the hydrophobic cluster in the allosteric site to stabilize helix 3 in the R-state, while the smallest side chain at R97 (i.e. R97A) does not pack at all and leaves helix 3 shifted toward switch II in the T-state. Increasing helix 3 flexibility by removing the R97 side chain altogether (i.e. R97G) appears to decrease the barrier between states, resulting in a mixture of the R– and T-states.

### Allosteric site mutants stabilize the R-state through compensatory interactions

We next focused on the details of packing in the allosteric site to understand the mechanism through which each R97 mutant favors the active site in the R-state or T-state. In our canonical R-state structure of HRAS, Ca^2+^ and acetate bind in the allosteric site to stabilize helix 3 toward helix 4, with backbone interactions between Y137 of helix 4, R97 in helix 3, and D107 of loop 7 (**see inset**, **Figure 3A**). Although Ca(AOc)_2_ is present in the crystallization mother liquor in all R32 crystals, calcium and acetate did not bind in the allosteric site in any of the structures and instead we observed a single water molecule in each of the R-state favoring structures (**‘bridge water’,** **Figure 3A**). This water molecule forms H-bonds with the backbone carbonyl groups of both Y137 and D107, replacing the interactions observed for Ca^2+^ in the wild type HRAS structure. Simultaneously, the side chain of Y137 shifts slightly to optimize packing with the various hydrophobic side chains at residue 97. Furthermore, K101 in R97V, R97L, and R97M was shifted toward the solvent to overlap with the position occupied by R97 in the wild type HRAS in the R-state. In the R-state mutants, the conformation of K101 is stabilized by proximity to D107 (**Figure 3A**). Thus, in addition to the water molecule between helix 4 and loop 7, the interaction between the side chains of D107 and K101 appears to compensate for the loss of the acetate-R97 interaction. R97I is an exception, as K101 is disordered in that structure due to D107 taking on an alternate conformation (**magenta structure**, **Figure 3A**).

**Figure 3:**
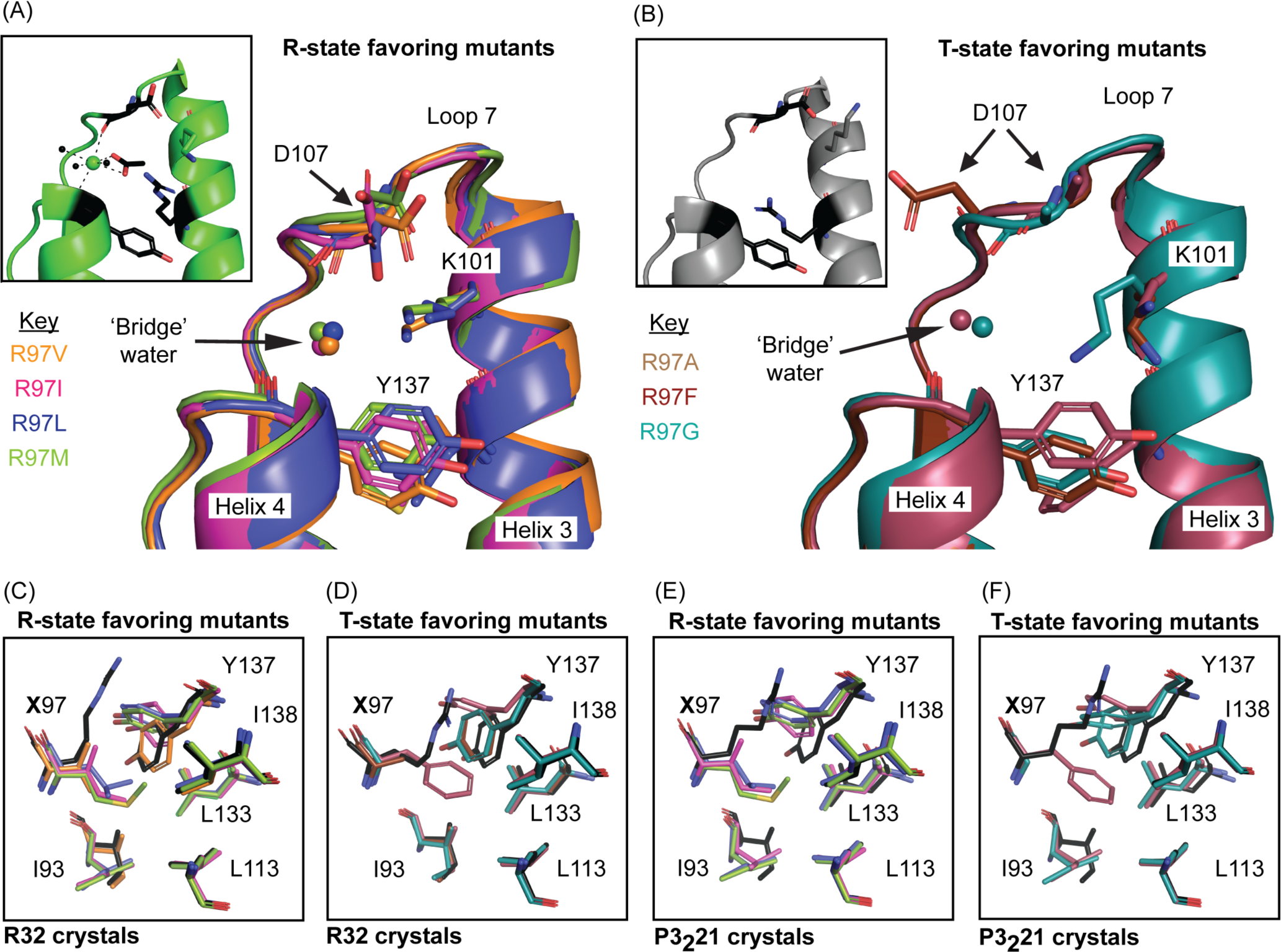
R97 mutants promote compensatory interactions in the allosteric site and hydrophobic pocket. (A) R-state favoring mutants show a ‘bridge’ water in the allosteric site that replaces Ca^2+^ while K101 compensates for loss of the guanidinium group of R97. Inset shows the allosteric site of wildtype HRAS in the R-state (PDB code 3K8Y ^13^). Y137, R97, K101, D107, acetate and water molecules coordinated to Ca^2+^ (green) are shown in black. (B) With the exception of R97G, the T-state favoring mutants do not show compensation by K101, nor closure of the allosteric site. Inset shows the allosteric site of wildtype HRAS in the T-state (PDB code 2RGE ^21^), with Y137, D107, R97, and K101 in black for reference. (C) and (D) are the hydrophobic pockets of R97 mutants crystallized in the R32 space group. (E) and (F) are the mutants crystallized in the P3_2_21 space group, with wildtype HRAS shown in black for reference (PDB code 1CTQ ^22^).

Consistent with our active site observations, loop 7 and helix 3 in the T-state favoring mutants R97A and R97F were shifted away from helix 4 (**Figure 2C** and **Figure 3B**). However, the allosteric site water molecule was present in the crystal structure of R97F (**Figure 3B**), indicating that binding of this water molecule is not sufficient to drive the T– to R-state shift. As expected, K101 in R97A and R97F is oriented away from the allosteric site, as it is in the wild type HRAS in the absence of Ca(AOc)_2_ (**see inset,** **Figure 3B**), suggesting that this position is a characteristic of the T-state. In contrast, R97G favored K101 placement into the allosteric site, and overall showed allosteric site features like the R– state favoring mutants (**cyan structures,** **Figure 2C** and **Figure 3B**).

The overall residue conformations in the hydrophobic pocket for the R97 variants remain similar to those observed in the wild type structures, as they are likely to be important for the stability of the protein core. However, conformational changes do occur for residues I93 and Y137 adjacent to the mutation site (**Figure 3C-D**). In both the R– and T-states of wild type HRAS, the tyrosyl ring of Y137 interacts with the hydrocarbon moiety of R97 (**see inset,** **Figure 3A-B**). In R97M, R97L, and R97I, Y137 also packs against the mutant 97 residue, but is rotated 90° (**Figure 3C**). Furthermore, the I93 side chain in these structures is found in a different rotamer thereby making room for the bulky hydrophobic side chains at residue 97. The smaller V97 side chain preserved the conformation of both Y137 and I93 as in the wild type structure. Notably, the rotated tyrosyl group of Y137 H-bonds with helix 3 residues H94 and E98 in the R-state mutants (*not shown*), while this is not observed for the T-state favoring mutants (*below*). Though subtle, these compensatory packing alterations demonstrate the influence of the hydrophobic cluster at the base of the allosteric site on the global conformation of HRAS. Likewise, since the R97V structure revealed minimal changes in allosteric site interactions, its conformation of switch II and helix 3 were the most consistent with wild type HRAS in the R-state (**Figure 2C**).

In R97F, R97G, and R97A, rotation of Y137 is only seen in the presence of F97, while the conformation of I93 was unaltered in each of these structures (**Figure 3D**). Thus, changes in the conformation of I93 and Y137 correlate to some degree with the R– and T-states. However, the nature of their influence on this global transition was not immediately obvious. Coincidentally, the HRAS R97 mutants, except for R97V and R97A, also crystallized with P3_2_21 symmetry in the same conditions that yielded crystals with R32 symmetry (**Supplementary Table 3**). Unlike R32 crystals of HRAS, P3_2_21 crystals show more extensive crystal contacts at the allosteric site, helix 3 and switch II ^22;23^. Furthermore, helix 3 in this crystal form is stabilized in the R-state conformation due to crystal packing. Thus, P3_2_21 crystals provide a view of the allosteric effects of the R-state conformation on I93 and Y137, not only in the R-state mutants, but also on those that are found in the T-state in R32 crystals where switch II is free of crystal contacts. As expected, we observed the same series of packing interactions for the R-state mutants in the P3_2_21 crystals as were seen for their R32 crystals. This is because the observed changes in I93 and Y137 are necessary to accommodate the bulky side chains at residue 97 within the more closely packed allosteric site associated with the R-state (**Figure 3E**). Interestingly, while the R97A, R97G, and R97F mutants retain their I93 side chain conformations as observed in the wild type HRAS structures in their R32 crystals, both the bulky R97F side chain and non-side chain mutant R97G in P3_2_21 crystals show I93 side chain conformations as in the R-state mutants (**Figure 3F**). While Y137 is near the allosteric site surface with room to adjust its conformation, the I93 residue is packed in the protein core and is located on helix 3 opposite the active site. Thus, the R-state promoted by crystal packing requires adjustment of I93 for bulky side chain mutations of R97. Conversely, changes in the I93 side chain conformation could affect the active site with a potential impact on GTP hydrolysis.

A role of I93 in the global Ras dynamics has been suggested before from ^15^N NMR experiments ^24;25^, consistent with our structural observations of changes in conformations of this side chain in response to mutation of R97. Likewise, Y137 is more dynamic than suggested by the crystal structures alone, as it is a site of phosphorylation in HRAS by ABL kinase ^26^. Changes in the conformations of I93 and Y137 revealed in P3_2_21 crystals of R97G suggest increased conformational plasticity in the allosteric site in the absence of a residue 97 side chain. In these crystals, I93 is in its alternate rotamer, despite no obvious cause due to packing, and Y137 is in the alternate rotated conformation (**Figure 3F**), as observed only for the R-state mutants in R32 crystals. Overall, it appears that Y137 and I93 are connected to the active site in a manner that correlates with the R-state and T-state conformations. R– state mutants require a change in packing at the adjacent hydrophobic cluster, while T-state mutants accommodate the hydrophobic cluster as in the wild type due to a more spacious allosteric site with helix 3 shifted toward switch II. In crystals where the active site is stabilized in an R-state like conformation with helix 3 shifted toward helix 4 (i.e. P3_2_21 crystals) the conformational changes in I93 and Y137 are observed to accommodate this shift in the context of the bulky mutant side chains as expected. However, adjustment of I93 in response to active site R-state stabilization without an obvious cause as seen in the R97G mutant suggests that I93 participates in the communication between the allosteric and active sites in a way that is not fully understood.

Under our experimental conditions, neither R97V nor R97A crystalized in the P3_2_21 space group. In the case of R97V, this may be due to this mutant favoring an active site *most* similar to wild type HRAS in the R-state (**Figure 2C**). In contrast, R97A stabilizes ordering of switch II into the anti-catalytic T-state conformation, which is inconsistent with crystal packing of P3_2_21 (**Figure 2C**). Given the small side chain of residue A97, it was expected to behave like the R97G mutant, with electron density maps showing a mixture of R– and T-state conformations. However, crystals of HRAS R97A grown at 18°C yielded a structure with full occupancy of the anti-catalytic T-state conformation without calcium or acetate bound in the allosteric site (**Figure 2** and **Figure 3**). Interestingly, we did observe the R-state when crystals of HRAS R97A were grown under the same solution conditions but at 4°C (**Figure 4**). It appears that a decrease in crystal growth temperature stabilizes the R97A R-state, with switch II making extensive intramolecular contacts in a more compact structure. At 4°C, R97A adopts a more packed allosteric site, with the water molecule present and both K101 and helix 3 shifted toward the site (**Figure 4A**). As expected, the conformations of I93 and Y137 were unchanged in this structure, as the alanine side chain allows for the wild type conformations associated with the hydrophobic cluster adjacent to the allosteric site. Likewise, Y96 was shifted toward switch II, and switch II was fully ordered in an R– state conformation (**Figure 4A**). The active site was also stabilized in the R-state (**Figure 4B**), as observed in all the R-state favoring R97 mutants (**Figure 2A**). It is not surprising that a mutation at R97 which has a propensity for both R– and T-state conformations, would favor the more packed R-state structure at the lower crystal growth temperature and the more disordered T-state at room temperature, given increased thermal motions.

**Figure 4:**
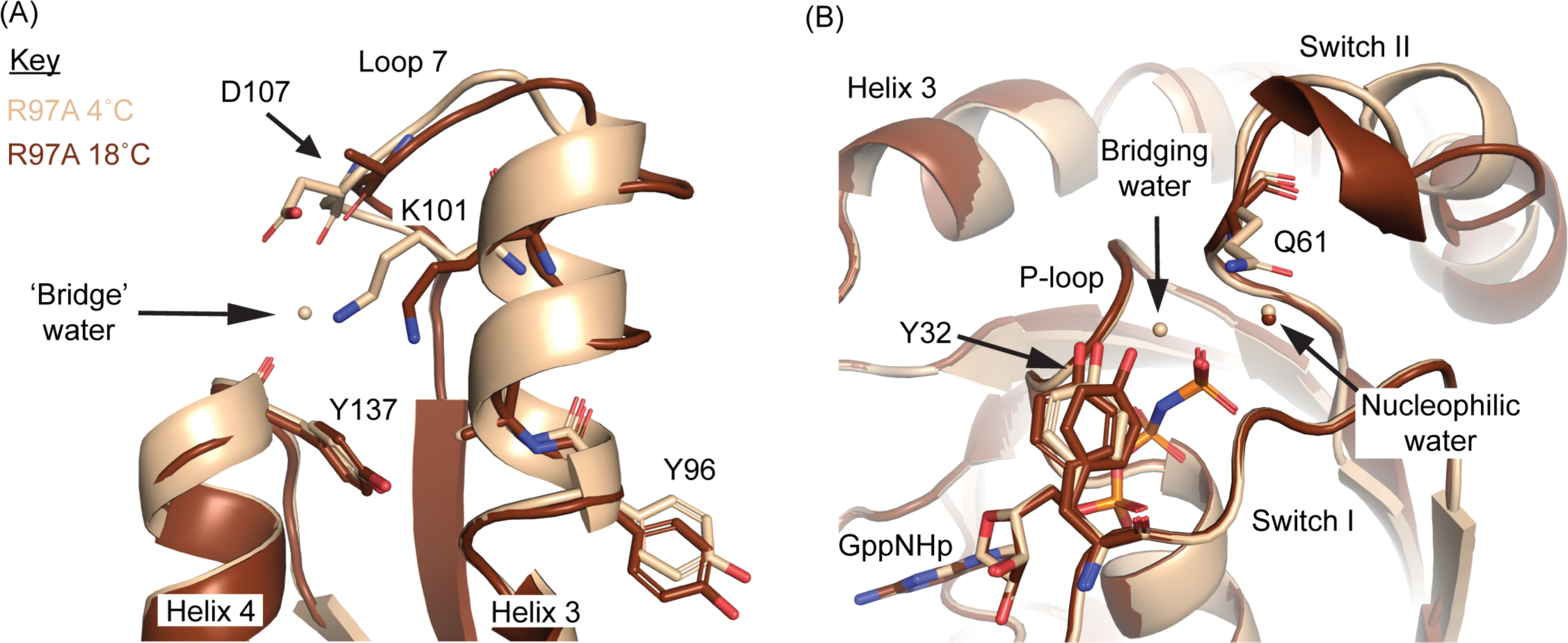
R97A crystals stabilize the R-state when grown at 4°C. (A) Crystals of R97A grown at 4°C (beige) stabilize the allosteric site in the R-state conformation (B) 4°C crystals of R97A show an active site consistent with the R-state conformation.

### The R-state enhances GTP hydrolysis by HRAS

Our experiments indicate that the transition between R– and T-states is dynamic and that the balance of conformational states between them can vary both with point mutations and with temperature. To correlate these conformational shifts with biochemical outcomes at physiologically relevant temperature, we performed single-turnover GTP hydrolysis assays at 37°C (**Supplementary Table 4**) and compared the first-order rate constants of these reactions to allosteric site packing in each of the variants. Since the placement of Y137 and M111 hardly shifts within our current structures, and R102 moves with helix 3, we used Heron’s rule to calculate the triangular area between the C_α_s of Y137, M111 and R102 in the R32 crystal structures as a way of measuring the allosteric site closure indicative of tighter packing between helices 3 and 4.

Comparisons between allosteric site closure and GTP hydrolysis are shown as orange dots in **Figure 5A**. Closure of the allosteric site correlates with an increase in the intrinsic hydrolysis rate constant of GTP, reflecting an increase in R-state populated HRAS as observed in our crystal structures of the R-state R97 mutants (**R97V, R97L, R97I, and R97M in** **Figure 5A**). Even R97G, which favored R-state features outside the active site, showed an increase in its hydrolysis rate constant (**orange circle,** **Figure 5A**). Likewise, the T-state favoring mutants R97F and R97A showed lower rate constants than wild type HRAS. For R97A, these hydrolysis data were consistent with the observed T-state conformation at the higher crystallization temperature (**red circle,** **Figure 5A**).

**Figure 5:**
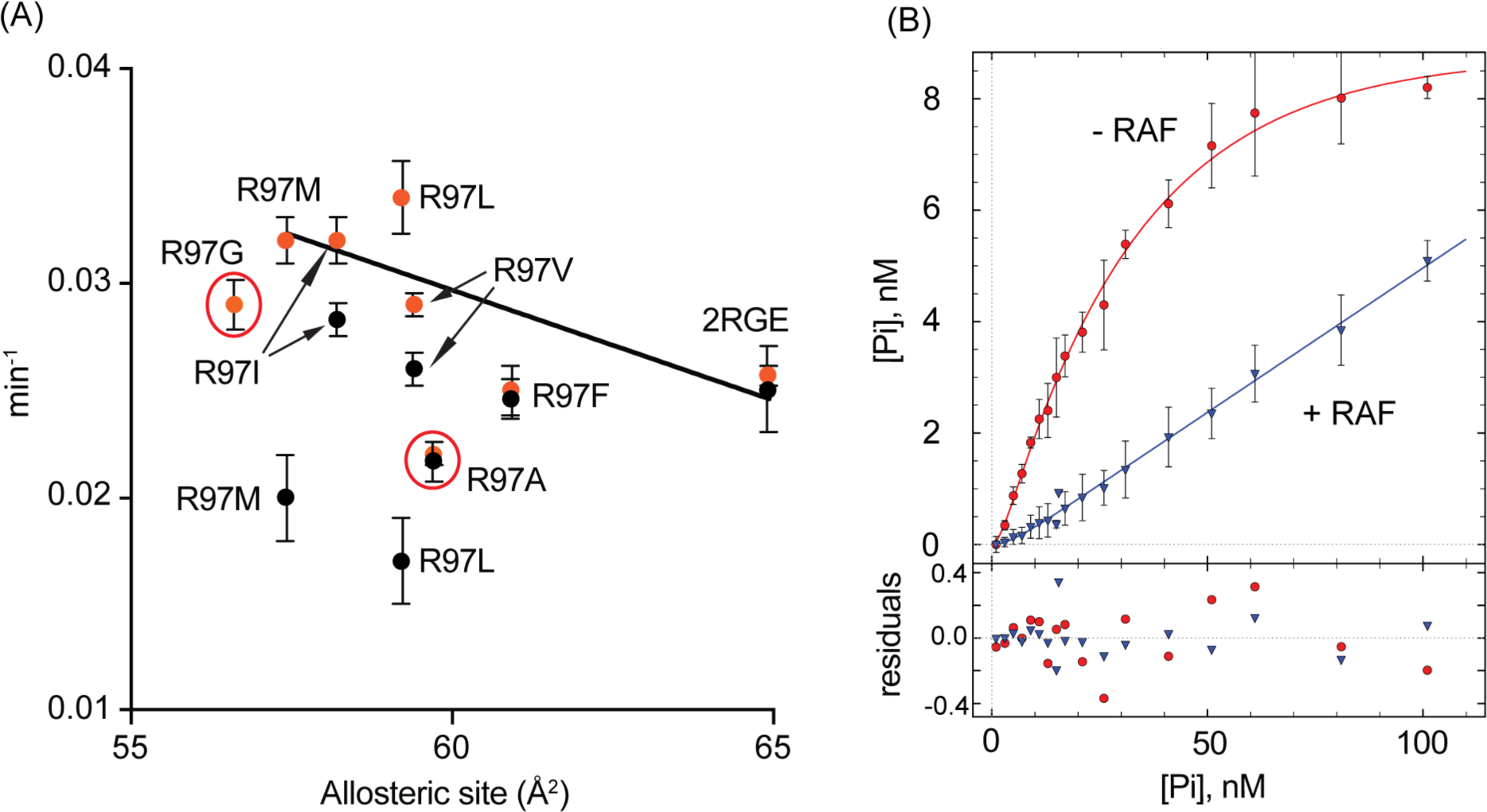
GTP hydrolysis is sensitive to both allosteric site mutation and binding to RAF1-RBD. (A) Correlation of allosteric site closure and intrinsic hydrolysis. Orange dots reflect first-order rate constants for intrinsic GTP hydrolysis of the R97 mutants. Black dots also reflect first-order rate constants, but in the presence of excess purified RAF1-RBD. (B) Single-turnover reaction for HRAS R97G alone (red line) or in the presence of excess RAF1-RBD (blue line). Each dot represents at least triplicate measurements of radioactive P_i_ at the indicated time point.

Part of the allosteric model for intrinsic hydrolysis involves stabilization of switch I in a closed conformation by the RAS effector RAF (**step 1,** **Figure 1A**). Since RAF binds switch I but not switch II, the T-state mutants are expected to retain a disordered switch II and RAF binding should not alter GTP hydrolysis, as is the case for wild type HRAS (2RGE in **Supplementary Table 4**). Indeed, binding of RAF1-RBD had no effect on GTP hydrolysis for the T-state mutants (e.g. R97F and R97A) (**Supplementary Table 5**). However, our model predicts that addition of RAF1-RBD should synergize with R-state favoring mutants of R97, stabilizing switch I and switch II to enhance GTP hydrolysis. However, the hydrolysis rate constant was reduced to varying degrees in the R-state mutants (**black dots,** **Figure 5A**). Of special note was GTP hydrolysis by R97G in the presence of RAF1-RBD, which was too slow to model in any significant way (**Figure 5B**). Clearly, the ordering of switch II in the presence of RAF1-RBD with a stabilized switch I is not sufficient to promote GTP hydrolysis. Perhaps the alternative conformations of I93 and Y137 we observe in our R97 mutants help to decouple the cooperative motions that occur between the allosteric site, helix 3, and switch II, particularly regarding the placement of catalytic residue Q61 for hydrolysis of GTP.

Taken together, our data show unequivocally that the allosteric site is in communication with the active site of HRAS, where it influences the rate of GTP hydrolysis on HRAS. However, GTP hydrolysis is impaired by the combined effect of RAF binding, and the conformational changes at I93 and Y137 necessary for packing of the R97 mutants’ bulky side chains. This is demonstrated in the decrease in GTP hydrolysis of R97G and the R-state favoring mutants, particularly for those with normal or slightly enhanced GTP hydrolysis in the absence of RAF1-RBD. Binding of RAF1-RBD quenches switch I dynamics in HRAS while simultaneously increasing motions in switch II ^27;28^. This could alter catalytically productive placement of Q61 in the active site for the allosteric site mutants in the presence of RAF1-RBD.

## Conclusions

Here, we establish a functional link between the active and allosteric sites in HRAS by showing that modulation of the allosteric site by mutagenesis affects the rate constants for GTP hydrolysis. This link was previously proposed based on structural analysis of HRAS in a crystal form where switch I is in the conformation observed in the RAS/RAF1-RBD complex and switch II is free of crystal contacts (PDB ID 3K8Y) ^13^. An H-bonding network linking the allosteric site to the active site was clearly observed, leading us to predict that measuring hydrolysis rate constants in the presence of Ca(AcO)_2_ should reveal enhanced intrinsic hydrolysis of GTP on RAS, given that the binding of calcium and acetate to the allosteric site was expected to order the active site in solution as it does in the crystals. This turned out not to be the case, consistent with NMR experiments that aimed at probing the metal binding properties of the allosteric site in solution. The NMR experiments revealed weak and non-selective binding of metal ions ^16^, contrary to what was observed in the crystals, where Ca^2+^ was clearly selected over Mg^2+^ ^13^. Since these early experiments, we have shown that RAF1-RBD promotes robust dimerization of RAS on supported lipid bilayers and to a smaller extent in solution ^29^. The crystal structure of the HRAS/RAF1-RBD complex (PDB ID 4G0N) shows that RAS forms a dimer through helices α4 and α5 (the α4– α5 dimer), generated through a 2-fold crystallographic symmetry axis ^28^. This dimer appeared again in a structure of KRAS/RAF-RBD-CRD (PDB ID 6XI7) ^30^. The α4– α5 dimer is also present in crystals of HRAS (PDB ID 3K8Y) where we first observed the connection between the allosteric and active sites ^13^. As the allosteric site is adjacent to the dimer interface, it is possible that the Ca^2+^ binding site is stabilized in the RAS dimer, providing increased affinity and specificity for Ca^2+^. This would explain the solution NMR experiments under conditions where RAS is entirely in its monomeric form ^16;31^ and the lack of response to Ca^2+^ in terms of GTP hydrolysis rate constants in monomeric RAS. As the Ca^2+^ binding issue remains unresolved, site directed mutagenesis was used in the current study to systematically perturb the allosteric site and show that the proposed R-state modulated by a shift in helix 3 toward helix 4 indeed correlates with hydrolysis of GTP on RAS.

The results presented here suggest two critical points that are generally relevant to small GTPases. First, more attention to allostery is needed to properly understand the intrinsic GTP hydrolysis reaction performed by different small GTPases. While we focused on HRAS here, the role of allostery in regulating the function of oncogenic mutants of NRAS, and even more so KRAS, are also needed. These two isoforms are well conserved between HRAS (95% sequence similarity), yet some amino acid substitutions exist near the allosteric site and network ^12^. These isoforms differ from HRAS in their dynamics and ability to promote GTP hydrolysis ^12;15; 32-34^. Whether the allosteric site plays a role in these functional differences, and how the allosteric site interacts with oncogenic mutations in the active site, is a necessary next step to a better understand these critically important enzymes. Second, the study of small GTPase allostery will likely provide a more complete picture of their evolution as signaling proteins. RAS, as well as many other small GTPases, are unlike other well-studied enzymes in that they have evolved to have poor catalytic efficiency on their own ^35^, and thus could be greatly susceptible to allosteric modulation of GTP hydrolysis. For instance, despite structural similarities between ATPases and GTPases, such as the well conserved P-loop, enzymes of these two families can show up to six-orders of magnitude differences in their phosphoryl-transfer capabilities ^36;37^. While the experiments described here looked at mutations of a single residue, they demonstrate that the allosteric site could be an underappreciated region in this evolution.

While our experiments show a functional connection between the allosteric site and the active site, the role of RAF in our hydrolysis experiments shows more complexity than implied in our previously published mechanism of intrinsic hydrolysis ^13^, where RAF primarily played the role of stabilizing switch I in a closed conformation (**Figure 1**). At that point we had not yet considered the effect of binding RAF on increased dynamics of switch II ^28^. The experiments we present here show that binding of RAF to RAS alters the balance of R– and T-conformational states in the direction of the T-state, determined by a lower GTP hydrolysis rate constant. This effect varies among the allosteric site mutants. Thus, the increase in switch II dynamics seen for HRAS-RAF complexation represents a mechanism to destabilize the R-state ^27;28^, which appears to synergize with the allosteric site mutants in our study. This synergy appears to be important, as the dynamic change in switch II has little effect on intrinsic GTP hydrolysis for wild type HRAS in complex with RAF-RBD.

Interestingly, the work presented here provides an explanation for a cluster of poorly studied RAS mutations found in human tumors (**Figure 6**) ^38^. From our hydrolysis experiments with RAF1-RBD, we can infer that some of these mutations may inhibit GTP hydrolysis on the RAF1-RAS complex to enhance signaling through MAPK signaling pathway and thereby promote cell growth. For instance, it is likely that RAF1 will cooperate with NRAS R97G and KRAS R97I to decrease GTP hydrolysis. Some of the allosteric site oncogenic mutations may destabilize the R-state by impairing the cooperative interactions that occur between helix 3, loop7 and helix 4, or interrupting binding with a putative allosteric ligand, thereby increasing the overall activated state of RAS in the cell (**orange residues,** **Figure 6**). In the case of I93V, the valine mutation will essentially mimic the preferred rotamer of I93 in our R-state favoring structures of HRAS, indicating that it could influence the T-to R-state transition via destabilizing hydrophobic packing beneath the allosteric site. Other mutations found in this pocket may have similar effects (**yellow residues,** **Figure 6**). Thus, dissection of the allosteric site will provide a better understanding of disease mutations that have been difficult to characterize ^5^.

**Figure 6:**
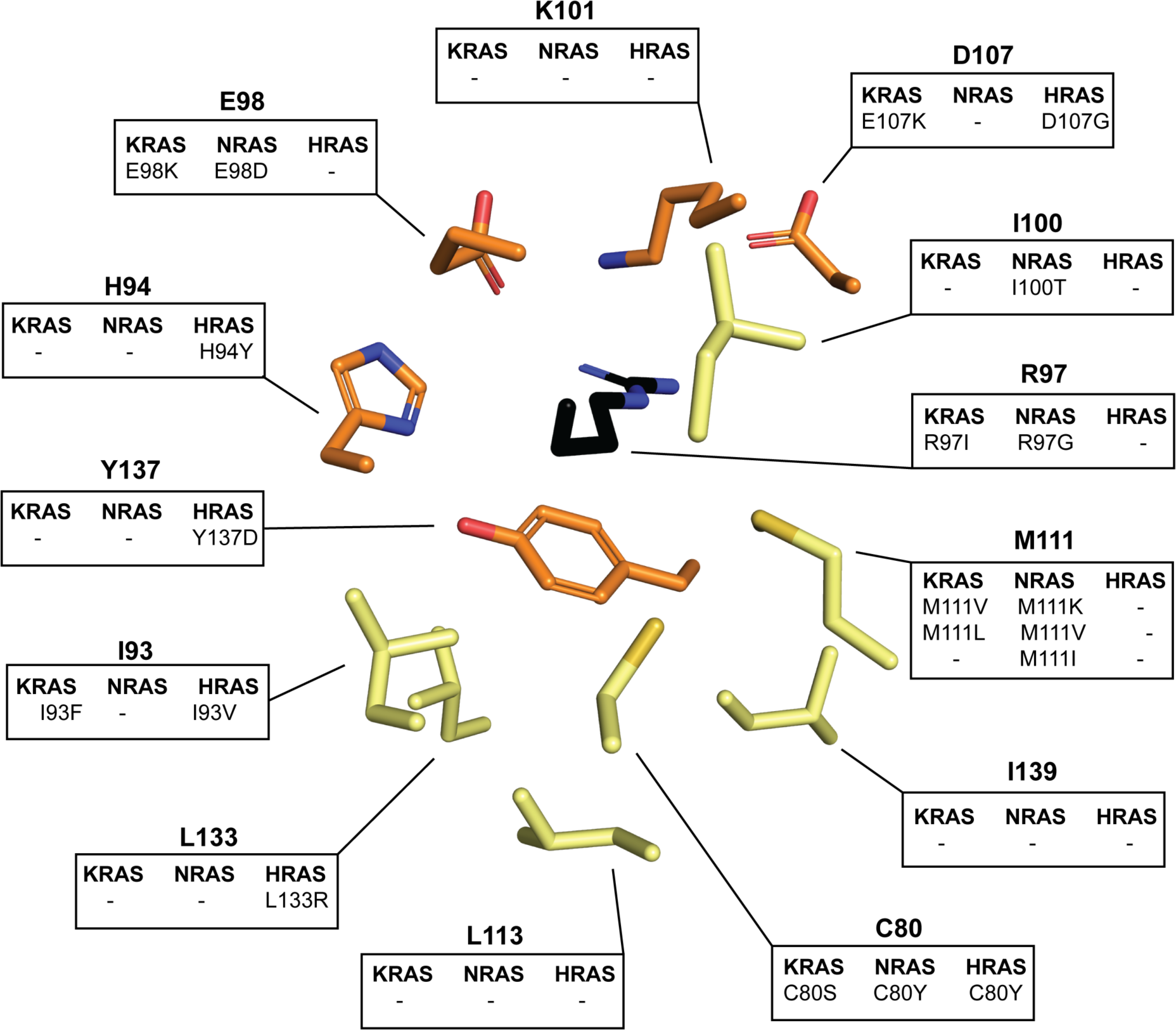
Mutations in the allosteric pocket found in cancer. Boxes represent mutations found at that residue. Residue colors are explained in the text. All mutations were identified in the COSMIC database.

Finally, this work revitalizes the possibility of utilizing intrinsic hydrolysis to attenuate or block the oncogenic activities of RAS. While the most common oncogenic mutations in KRAS target GAP-mediated hydrolysis, many (if not most) oncogenic mutations retain some form of intrinsic hydrolysis ^5^. Thus, targeting the allosteric site may be a potential means to activate intrinsic hydrolysis. Indeed, we previously showed that the allosteric site can be utilized to drive G12V mutants of HRAS into the R-state when crystals of this mutant are soaked in Ca(OAc)_2_ ^14^. More studies on the allosteric site are needed to understand its role in cellular homeostasis and to leverage it against RAS driven cancers.

## Materials and Methods

### Experimental design to explore allostery in Ras

The allosteric site consists of a ring of polar (Y137, H94) and charged residues (D107, K101, E98) surrounding R97, which is poised over a hydrophobic pocket formed by the sidechains of I93, C80, M111, L113, L133, Y137, I139 and the aliphatic carbon chain of R97 (**Figure 1B** **and** **Figure 6**). A potential space for packing in the hydrophobic pocket was observed during initial analyses and R97 was mutated to hydrophobic residues. This could drive the R-state transition by packing of the mutated side chain in the core, thereby shifting the helix 3 balance of conformational states toward helix 4. Moderate (i.e. R97V, R97L, R97I) and large (i.e. R97M, R97F) packing mutations were tested, as well as different side-chain configurations (i.e. R97L vs. R97I, R97M vs. R97F). Moderately sized groups were expected to favor the R-state and the larger hydrophobic groups to sterically prevent the motion of helix 3 toward helix 4, which is necessary for the R-state. Increased flexibility (R97G) and loss of packing (R97A) mutations were also tested. The large hydrophobic mutants were expected to favor the T-state due to steric interactions within the allosteric stie, and the R97A and R97G mutants were expected to favor the T-state due to increased dynamics of both helix 3 and switch II.

### Mutagenesis, protein purification and enzymology

All mutant protein for crystallization and hydrolysis used residues 1-166 comprising the catalytic G-domain of HRAS (EC 3.6.5.2). Residue 97 mutations to glycine, alanine, valine, isoleucine, methionine and phenylalanine of HRAS were made using a two-stage mutagenesis protocol ^39^, based on QuikChange parameters. The RAS proteins were purified as previously described ^40^ and so was C-RAF1-RBD (RAF1-RBD; EC 2.7.11.1) ^26^. In order to crystallize HRAS mutants in their active form (i.e. GTP bound), we exchanged GDP for a non-hydrolyzable GTP-analogue guanylyl-5-imidodiphosphate (GppNHp) using an established protocol ^40^. Protein for hydrolysis was kept bound to GDP, and both the GDP and the GppNHp bound proteins, were transferred into stabilization buffer (20 mM HEPES, 50 mM NaCl, 20 mM MgCl_2_, 1 mM DTT, pH 7.5). Immediately after buffer exchange, protein was flash frozen in liquid nitrogen, in 20-100µL aliquots, and stored at –80°C. Protein purity was determined using SDS-PAGE and concentration was determined to be 12-20 mg/mL using the Bradford assay ^41^.

Single turnover hydrolysis reactions for the R97 mutants were performed as previously published ^42^, with small modifications described here. All reactions were performed at 37°C. GDP-bound RAS (5µM) was preloaded with ^32^P-γ-GTP (50nM) from Perkin Elmer, for 5 minutes in exchange buffer (20 mM Tris, 1 mM EDTA, 2 mM DTT, pH 8.0) in a 100µL reaction. Once nucleotide was exchanged, initiation of hydrolysis was done by adding 4µL of nucleotide exchange reaction, to 16µL of pre-warmed hydrolysis buffer (20 mM Tris, 100 mM NaCl, 5 mM MgCl_2_, 2 mM DTT, and pH 8.0), to produce a 5-fold dilution of RAS and nucleotide concentration. Sixteen total reactions were performed for each single-turnover experiment, corresponding to 16 time points at 0, 2, 4, 6, 8, 10, 12, 14, 16, 20, 25, 30, 40, 50, 60, 80 and 100 minutes. Amount of ^32^P_i_ formed for each time point was determined by organic extraction ^43;44^, and detection of β-emission was done using a HIDEX liquid scintillation counter. Recorded CPM and TDCR measurements were used to determine the fmols of ^32^P_i_ formed during the reaction using a published procedure ^42^. Reactions were then converted into concentration (nM) of ^32^P_i_ before analysis.

### Kinetic Modeling

Dyanfit4 was used to determine the rate constants for intrinsic hydrolysis (*k*_hyd_) for each of the HRAS R97 mutants ^45;46^, using protocols that we previously published in detail for RAS proteins ^12^. For each reaction, two parameters were determined: *k*_hyd_ and the starting concentration of RAS-GTP. Fitting of RAS-GTP was performed to mitigate errors in the estimation of protein concentration, titration errors, and differences in GTP concentration due to radioactive decay. Since the association and dissociation rates for binding of RAF1-RBD to HRAS are significantly faster than the intrinsic hydrolysis of Ras ^47^, we assumed that the association and dissociation rates were also significantly higher for the HRAS mutants and so were not included in the parameter model. We were successful in using a first-order kinetic mechanism for wild type HRAS and all the allosteric site mutants except for R97G in the presence of RAF1-RBD, where the reaction was too slow to measure (**Figure 5B**). Rates and rate constants determined by Dynafit4 can be found in **Supplementary Tables 4** and **5**.

### Protein crystallization

All protein for these crystallization experiments were prepared in the same way as the protein used for our hydrolysis assays. Crystallization was performed using the sitting drop and vapor diffusion methods using various concentrations of Ca(OAc)_2_ and PEG 3350 for each mutant around the original hit condition 28 from the Hampton Research PEG Ion Screen. We obtained crystals of HRAS R97G, R97I, R97L, R97M, and R97F mutants with symmetries of both R32 and P3_2_21 space groups, while the R97A and R97V mutants were only crystallized with symmetry of the R32 space group. The crystals were grown in 2µL by 2µL protein to mother liquor drops. Prior to X-ray diffraction and data collection, crystals were briefly soaked in mother liquor containing 30% glycerol for cryoprotection. X-ray diffraction data for R97F crystallized with P3_2_21 symmetry, and all of the HRAS R97G, R97A, R97V, R97I, R97L, and R97M bound to GppNHp data were collected on a home source instrument (Rigaku MicroMax 007 R-AxisIV^++^). Data for the R97F and R97A crystals with R32 symmetry were collected on the ID-22 SER-CAT beamline at the Advanced Photon Source (Argonne National Laboratory). Diffraction data were collected at 100K for both the home and synchrotron X-ray sources.

Indexing, integration, scaling and post-refinement were performed on HKL3000 ^48^, and data collection statistics can be found in **Supplementary Table 1 and 3**. Molecular replacement was performed using the default settings in the auto-MR program in the PHENIX suite of programs ^49^; the PDB model 1CTQ was used as a phasing model for the P3_2_21 crystals, and PDB model 3K8Y was used as a phasing model for the R32 crystals—ligands were not included in molecular replacement. After molecular replacement, starting models were edited for the correct mutation, and ligands (i.e., GppNHp-Mg), were set to full occupancy. PHENIX was then used to perform a single round of simulated annealing to remove model bias. Further refinement and model building was performed by alternating use of PHENIX and the model-building program COOT ^50^. Refinement statistics can be found in **Supplementary Table 1 and 3**.

**Supplementary material description:**

Table 1: Statistics R32 protein crystals

Table 2: Crystallization conditions

Table 3: Statistics P3_2_21 protein crystals

Table 4: Hydrolysis rates and rate constants for H-Ras R97 mutants

Table 5: Hydrolysis rates and rate constants for H-Ras R97 mutants in the presences of Raf1-RBD

## Acknowledgements

This project was originally funded by NSF MCB 1244203 and completed with NSF MCB 2121426 awarded to Carla Mattos.

## Contributions

Carla Mattos conceived of the project. Susan Fetics crystallized and solved the structure of R97L in the P3_2_21 and R32 space groups. Kathleen Davis crystallized the R97A mutant, in both R-and T-state R32 space groups. Jose Rodrigues crystallized the R97G mutant in the R32 space group. Christian Johnson crystallized the remaining allosteric site mutants, designed experiments, performed all GTP hydrolysis experiments. Carla Mattos and Christian Johnson designed experiments and wrote the paper.

## Figure Legends

