## Supplemental Tables for "Allosteric site variants affect GTP hydrolysis on RAS"

**Supplementary Table 1: Crystal statistics for R32 protein crystals**

| | **R97F** | **R97M** | **R97L** | **R97I** | **R97V** | **R97G** |
|---|---|---|---|---|---|---|
| PDB accession code | 8ELK | 8ELY | 8FG4 | 8ELX | 8EM0 | 8ELU |
| **Data collection and processing** | | | | | | |
| Resolution range | 44.1 – 1.5 (1.6 – 1.5) | 45.3 – 2.0 (2.0 – 1.96) | 27.0 – 1.8 (1.9 – 1.8) | 33.5 – 2.0 (2.0 – 1.97) | 31.0 – 2.1 (2.2 – 2.1) | 36.9 – 1.9 (2.0 – 1.9) |
| Space group | R32 | R32 | R32 | R32 | R32 | R32 |
| **Unit cell** a, b, c (Å) α, β, γ (°) | 88.1 88.1 133.3 90 90 120 | 89.0 89.0 135.9 90 90 120 | 89.9 89.9 136.3 90 90 120 | 89.2 89.2 135.2 90 90 120 | 89.1 89.1 135.5 90 90 120 | 88.6 88.6 135.3 90 90 120 |
| Total reflections | 136194 (6340) | 149776 (7061) | 153321 (9281) | 146826 (14810) | 117281 (4396) | 1675401 (16338) |
| Unique reflections | 31673 (2882) | 14971 (1358) | 17828 (1289) | 14255 (1424) | 11612 (666) | 15658 (1556) |
| Multiplicity | 4.3 (2.2) | 10 (5.2) | 8.6 (7.2) | 10.3 (10.4) | 10.1 (6.6) | 10.7 (10.5) |
| Completeness (%) | 94.9 (73.6) | 99.2 (91.8) | 96.2 (70.4) | 94.6 (98.3) | 95.1 (56.1) | 99.7 (99.4) |
| Mean I/sigma (I) | 10.4 (0.5) | 13.9 (1.4) | 31.3 (3.7) | 17.1 (1.9) | 17.6 (1.6) | 22.8 (3.1) |
| Wilson B-factor | 20.4 | 26.6 | 32.9 | 22 | 25.5 | 26.5 |
| R-merge | 0.141 | 0.111 | 0.203 | 0.101 | 0.126 | 0.075 |
| **Structure solution and refinement** | | | | | | |
| R-work (%) | 17.9 (37.5) | 15.7 (26.9) | 18.6 (41.1) | 22.7 (43.3) | 15.6 (26.7) | 17.7 (25.8) |
| R-free (%) | 20.7 (41.7) | 20.2 (30.0) | 22.2 (36.2) | 28.6 (41.7) | 20.1 (33.2) | 20.9 (29.6) |
| **Atoms** | | | | | | |
| Macromolecules | 1309 | 1330 | 1321 | 1311 | 1312 | 1276 |
| Ligands | 37 | 43 | 35 | 35 | 35 | 35 |
| Solvent | 163 | 166 | 144 | 144 | 137 | 143 |
| **RMSD** | | | | | | |
| bonds (Å) | 0.007 | 0.008 | 0.009 | 0.008 | 0.008 | 0.007 |
| angles (˚) | 1.21 | 1.18 | 1.21 | 1.3 | 1.2 | 1.2 |
| **Ramachandran (%)** | | | | | | |

| | | | | | | |
|---|---|---|---|---|---|---|
| favored<br>allowed<br>outliers | 96.9<br>3.1<br>0 | 97.6<br>2.4<br>0 | 98.8<br>1.2<br>0 | 98.2<br>1.8<br>0 | 97.6<br>2.4<br>0 | 97<br>3.1<br>0 |
| Clashscore | 5.7 | 4.5 | 4.1 | 5.3 | 4.6 | 4.3 |
| **B-factor (Å$^2$)** | | | | | | |
| Average | 26.9 | 28.6 | 36.7 | 22.1 | 31.5 | 31.5 |
| Macromolecules | 25.9 | 27.5 | 36.3 | 21.8 | 31.2 | 30.9 |
| Ligands | 18.5 | 27 | 27.4 | 15.0 | 23.6 | 21.9 |
| Solvent | 36.7 | 37.2 | 42.4 | 27.4 | 36.7 | 39.5 |

**Supplementary Table 1 continued**

| | R97A (R-state) | R97A (T-state) |
|---|---|---|
| PDB accession code | 8ELS | 8ELW |
| **Data collection and processing** | | |
| Resolution range | 25.7 – 2.3 (2.3 – 2.27) | 30.7 – 1.7 (1.8 – 1.7) |
| Space group | R32 | R32 |
| **Unit cell** | | |
| a, b, c (Å) | 88.9 88.9 135.0 | 88.2 88.2 133.8 |
| α, β, γ (°) | 90 90 120 | 90 90 120 |
| Total reflections | 100301 (6016) | 243353 (24222) |
| Unique reflections | 9738 (955) | 22326 (2202) |
| Multiplicity | 10.3 (6.3) | 10.9 (11.0) |
| Completeness (%) | 99.73 (99.06) | 99.9 (100) |
| Mean I/sigma (I) | 16.2 (1.5) | 59.3 (13.8) |
| Wilson B-factor | 30.48 | 16.8 |
| R-merge | 0.143 | 0.06 |
| **Structure solution and refinement** | | |
| R-work (%) | 16.2 (23.9) | 16.0 (16.7) |
| R-free (%) | 21.2 (30.7) | 18.6 (20.2) |
| **Atoms** | | |
| Macromolecules | 1333 | 1361 |
| Ligands | 35 | 36 |
| Solvent | 110 | 148 |
| **RMSD** | | |
| bonds (Å) | 0.008 | 0.007 |
| angles (˚) | 1.1 | 1.1 |
| **Ramachandran (%)** | | |
| favored | 97.6 | 95.7 |
| allowed | 2.4 | 3.7 |
| outliers | 0 | 0.6 |

| | | |
|---|---|---|
| Clashscore | 4.1 | 3.3 |
| **B-factor (Å$^2$)** | | |
| Average | 35.1 | 22.4 |
| Macromolecules | 34.9 | 21.7 |
| Ligands | 26.6 | 14.5 |
| Solvent | 39.7 | 31.2 |

**Supplementary Table 2: Crystallization conditions for H-Ras R97 mutants**

| Mutant | PDB code | Crystal form | State | ˚C | pH | Ca(AOc) (mM) | PEG 3350 (%) |
|---|---|---|---|---|---|---|---|
| - | *3K8Y | R32 | R | 18 | 7.5 | 200 | 20 |
| R97G | | R32 | R | 18 | 7.5 | 145 | 24 |
| R97A | | R32 | R | 4 | 7.5 | 365 | ~20 |
| R97A | | R32 | T | 18 | 7.5 | 365 | ~20 |
| R97V | | R32 | R | 18 | 7.5 | 152 | 15 |
| R97I | | R32 | R | 18 | 6 | 133 | 18 |
| R97L | | R32 | R | 18 | 7.5 | 350 | 24 |
| R97M | | R32 | R | 18 | 7.5 | 152 | 25 |
| R97F | | R32 | T | 18 | 7.5 | 152 | 25 |
| R97G | | P3221 | - | 18 | 7.5 | 160 | 24 |
| R97I | | P3221 | - | 18 | 7.5 | 168 | 17 |
| R97L | | P3221 | - | 18 | 7.5 | 275 | 21 |
| R97M | | P3221 | - | 18 | 7.5 | 152 | 20 |
| R97F | | P3221 | - | 18 | 7.5 | 268 | 28 |

*Originally published in Buhrman et al. 2010

**Supplementary Table 3: Crystal statistics for $P3_221$ protein crystals**

| | **R97F** | **R97M** | **R97L** | **R97I** | **R97G** |
|---|---|---|---|---|---|
| PDB accession code | 8ELR | 8ELZ | 8FG3 | 8ELV | 8ELT |
| **Data collection and processing** | | | | | |
| Resolution range | 34.0 – 2.0 (2.1 – 2.0) | 34.0 – 1.7 (1.8 – 1.7) | 28.7 – 1.5 (1.5 – 1.49) | 33.8 – 2.2 (2.2 – 2.2) | 33.7 – 1.7 (1.7 – 1.66) |
| Space group | $P3_221$ | $P3_221$ | $P3_221$ | $P3_221$ | $P3_221$ |
| **Unit cell** a, b, c (Å) α, β, γ (°) | 39.3 39.3 158.0 90 90 120 | 39.2 39.2 157.7 90 90 120 | 39.7 39.7 157.6 90 90 120 | 39.0 39.0 158.1 90 90 120 | 38.9 38.9 157.4 90 90 120 |
| Total reflections | 86621 (6290) | 100390 (2193) | 137166 (13129) | 59840 (546) | 159560 (10181) |
| Unique reflections | 9481 (925) | 13525 (645) | 24494 (2387) | 7480 (273) | 17157 (1669) |
| Multiplicity | 9.1 (6.8) | 7.4 (3.4) | 5.6 (5.5) | 8.0 (2.0) | 9.3 (6.1) |
| Completeness (%) | 99.3 (100.00) | 89.4 (43.8) | 99.9 (100.0) | 91.2 (34.9) | 99.3 (99.5) |
| Mean I/sigma (I) | 24.9 (4.8) | 38.2 (5.1) | 42.8 (12.8) | 15.3 (1.9) | 22.3 (1.8) |
| Wilson B-factor | 32.4 | 25 | 16 | 24.9 | 29 |
| R-merge | 0.051 | 0.079 | 0.05 | 0.089 | 0.055 |
| **Structure solution and refinement** | | | | | |
| R-work (%) | 18.8 (19.8) | 17.9 (26.8) | 20.2 (24.7) | 18.4 (26.1) | 19.8 (34.5) |
| R-free (%) | 24.4 (31.0) | 22.7 (34.7) | 24.5 (30.8) | 27.0 (28.6) | 23.9 (38.0) |
| **Atoms** | | | | | |
| Macromolecules | 1311 | 1322 | 1304 | 1301 | 1327 |
| Ligands | 35 | 36 | 33 | 35 | 36 |
| Solvent | 77 | 139 | 171 | 73 | 124 |
| **RMSD** | | | | | |
| bonds (Å) | 0.008 | 0.007 | 0.007 | 0.008 | 0.007 |
| angles (˚) | 1.24 | 1.18 | 1.27 | 1.2 | 1.2 |
| **Ramachandran (%)** | | | | | |

| | | | | | |
|---|---|---|---|---|---|
| favored | 97.6 | 98.8 | 97.6 | 95.7 | 98.9 |
| allowed | 2.4 | 1.2 | 1.8 | 3.05 | 1.2 |
| outliers | 0 | 0 | 0.6 | 1.2 | 0 |
| Clashscore | 5 | 4.1 | 6.1 | 6.9 | 5.2 |
| **B-factor (Å$^2$)** | | | | | |
| Average | 34.2 | 26.1 | 21.5 | 29.9 | 31.3 |
| Macromolecules | 34.1 | 25.5 | 20.5 | 30 | 30.8 |
| Ligands | 27.9 | 22.5 | 14.2 | 24.5 | 25.8 |
| Solvent | 39 | 33.2 | 31 | 31.3 | 38.2 |

**Supplementary Table 4: Rate and rate constants for H-Ras R97 mutants alone**

| | $k_{hyd}$ | SE | CV% | Low (a) | High(a) | Rate (sec) | SD | n |
|---|---|---|---|---|---|---|---|---|
| **Wild type** | 0.0257 | 0.0008 | 3.1 | 0.0241 | 0.0274 | 0.216 | 0.0438 | 11 |
| **R97G** | 0.029 | 0.002 | 6.9 | 0.026 | 0.032 | 0.249 | 0.0123 | 3 |
| **R97A** | 0.022 | 0.001 | 4.5 | 0.02 | 0.025 | 0.184 | 0.0072 | 3 |
| **R97V** | 0.029 | 0.001 | 3.4 | 0.027 | 0.031 | 0.245 | 0.0017 | 3 |
| **R97I** | 0.032 | 0.002 | 6.2 | 0.027 | 0.036 | 0.253 | 0.0199 | 3 |
| **R97L** | 0.034 | 0.003 | 8.8 | 0.029 | 0.039 | 0.284 | 0.0086 | 3 |
| **R97M** | 0.032 | 0.002 | 6.2 | 0.029 | 0.035 | 0.279 | 0.0130 | 3 |
| **R97F** | 0.025 | 0.002 | 8.0 | 0.022 | 0.029 | 0.238 | 0.0144 | 3 |

(a) 95% confidence interval ; (b) Rate constant could not be determined.

**Supplementary Table 5: Rate and rate constants for H-Ras R97 mutants with Raf1-RBD**

| | $k_{hyd}$ | SE | CV% | Low (a) | High(a) | Rate (sec) | SD | n |
|---|---|---|---|---|---|---|---|---|
| **Wild type** | 0.025 | 0.002 | 8.0 | 0.022 | 0.028 | 0.263 | 0.0080 | 3 |
| **R97G(b)** | - | - | - | - | - | 0.049 | 0.0074 | 3 |
| **R97A** | 0.0217 | 0.0009 | 4.1 | 0.0199 | 0.0235 | 0.166 | 0.0066 | 3 |
| **R97V** | 0.026 | 0.0008 | 3.1 | 0.0244 | 0.0277 | 0.214 | 0.0073 | 3 |
| **R97I** | 0.0283 | 0.0008 | 2.8 | 0.0267 | 0.0299 | 0.230 | 0.0011 | 3 |
| **R97L** | 0.017 | 0.002 | 11.8 | 0.014 | 0.021 | 0.201 | 0.0257 | 3 |
| **R97M** | 0.02 | 0.002 | 10.0 | 0.016 | 0.024 | 0.141 | 0.0214 | 3 |
| **R97F** | 0.0246 | 0.001 | 4.1 | 0.0226 | 0.0266 | 0.210 | 0.0181 | 3 |

(a) 95% confidence interval ; (b) Rate constant could not be determined.
